# MidRISH: Unbiased harmonization of rotationally invariant harmonics of the diffusion signal

**DOI:** 10.1101/2023.08.12.553099

**Authors:** Nancy R. Newlin, Michael E. Kim, Praitayini Kanakaraj, Tianyuan Yao, Timothy Hohman, Kimberly R. Pechman, Lori L. Beason-Held, Susan M. Resnick, Derek Archer, Angela Jefferson, Bennett A. Landman, Daniel Moyer

## Abstract

**Objective:** Data harmonization is necessary for removing confounding effects in multi-site diffusion image analysis. One such harmonization method, LinearRISH, scales rotationally invariant spherical harmonic (RISH) features from one site (“target”) to the second (“reference”) to reduce confounding scanner effects. However, reference and target site designations are not arbitrary and resultant diffusion metrics (fractional anisotropy, mean diffusivity) are biased by this choice. In this work we propose MidRISH: rather than scaling reference RISH features to target RISH features, we project both sites to a mid-space.

**Methods:** We validate MidRISH with the following experiments: harmonizing scanner differences from 37 matched patients free of cognitive impairment, and harmonizing acquisition and study differences on 117 matched patients free of cognitive impairment.

**Conclusion:** MidRISH reduces bias of reference selection while preserving harmonization efficacy of LinearRISH.

**Significance:** Users should be cautious when performing LinearRISH harmonization. To select a reference site is to choose diffusion metric effect-size. Our proposed method eliminates the bias-inducing site selection step.

## I. Introduction

DIFFUSION weighted imaging (DWI) is a non-invasive, in-vivo magnetic resonance imaging modality used to understand white matter microstructure [1]. For each volume element, DWI captures the propensity for water to diffuse in a specific direction. Structural elements such as cell membranes and axonal fibers obstruct molecular movement, leading to higher observed diffusivity along these structures (parallel to the obstruction surfaces) in comparison to other directions incident to the structures [2]. Two metrics that characterize diffusion behavior are mean diffusivity (MD), which is the average magnitude of observed diffusion assuming a tensor (ellipsoidal) model, and fractional anisotropy (FA), a scaled variability measure of diffusivity, also assuming a tensor (ellipsoidal) model [2]. Studies have used DWI and related summary diffusion measures to study tissue changes caused by Alzheimer’s disease, schizophrenia, aging, multiple sclerosis and mild traumatic brain injury [3]–[10].

There are a growing number of multi-center studies that span multiple scanner, and acquisition protocols, or “sites”. Unfortunately, these site specifications introduce significant confounding differences in images and downstream analysis such as diffusion tensor imaging (DTI) metrics and connectomics [11]–[15]. Thus, there is a growing number of methods designed to reduce site-related differences whilst preserving biological variability, or “harmonization” [16], [17].

A popular harmonization method is the Linear Rotationally Invariant Spherical Harmonic method (LinearRISH), wherein diffusion signal is projected onto spherical harmonic basis functions [18], [19] and rotationally invariant power spectrum computed for each basis function set [20]. LinearRISH ascribes the site-wise differences to differences in these power spectra (the RISH coefficients). Previous works used LinearRISH harmonization for multi-site analysis of white matter abnormalities in schizophrenia [21] and connectomes of healthy traveling subjects [22]. Given a non-linear registration between all volumes, the LinearRISH procedure computes the shift needed to map each target site’s mean RISH coefficients to the mean RISH coefficients of a chosen reference site per template space voxel. After applying this procedure all sites should have the same average RISH coefficients as the reference site.

However, choosing a different reference site leads to different “harmonized” values. If a target site and the reference site were switched, the shift for each coefficient would change signs (Figure 1). This means the choice of reference site is non-trivial, and that, in a regression context like many of the use cases, choosing a reference site allows practitioners to choose effect size. At best ambiguity in this common procedure is not beneficial to accuracy or reproducibility, at worst it leaves a clear avenue for purposeful effect size inflation.

**Fig. 1.**
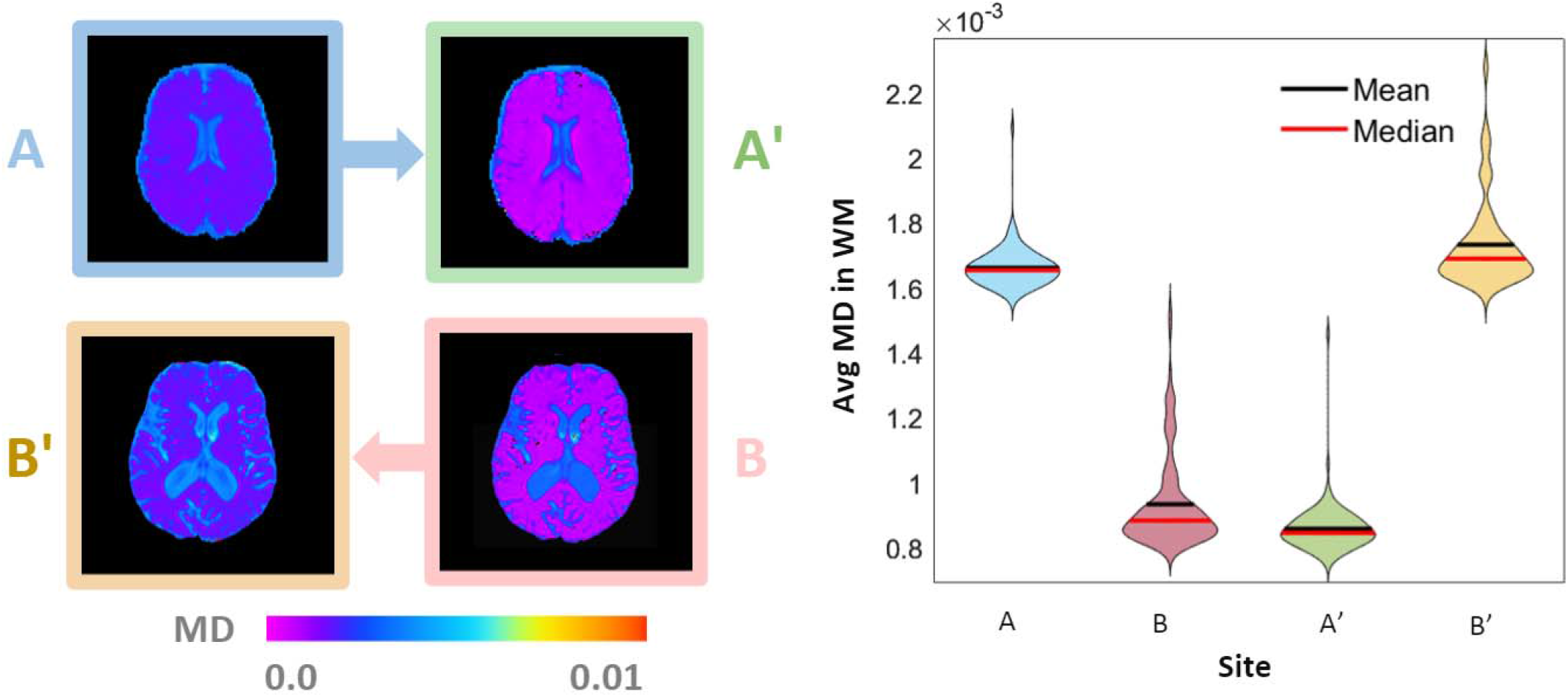
When site A (blue) is selected as the reference site for LinearRISH harmonization, site B’ (yellow) mean MD shifts up to the site A expected value. On the other hand, selecting site B (pink) as reference causes site A’ (green) to shift down to the site B expected value. It is up to the user to decide, which leads to arbitrary bias. Here we propose a quantitative solution.

In this paper, we propose a modification of the LinearRISH procedure to resolve this ambiguity. We propose shifting RISH coefficients to an unbiased midpoint between sites; this we show produces the mean of site-wise measurements in FA and MD.

## II. Methods

We demonstrate our method with two experimental designs. First, we compare MidRISH and LinearRISH performance at harmonizing data from the same study but different scanners. Then, we compare MidRISH and LinearRISH harmonization for data from two studies but same scanner specifications with b-value differences. Harmonization performance is based on site differences in mean MD and FA in cerebral white matter, spherical harmonic shape, and bundle-wise tractometry metrics.

### A. LinearRISH

Diffusion signal, 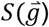, can be represented as a sum of products of spherical harmonic coefficients,*C*, and basis functions (Tournier), *Y* [18]. The sum is computed for all orders *l*, and degrees *m* where –*l* ≤ *m* ≤ *l*. We only consider even orders starting with 0 because odd orders are symmetric and sum to 0.

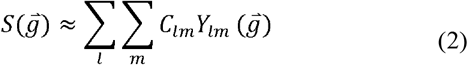

The RISH coefficients may be defined as

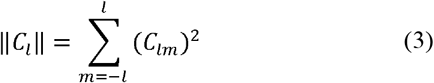

As their name suggests, these can be shown to be rotationally invariant (as the squared magnitude of each of the *l* orders is invariant under rotation). LinearRISH “harmonizes” these coefficients to be the same across sites. In order to do this, a diffeomorphic registration is computed (see D.4) and the expected coefficient value for each registered template voxel computed, denoted *E*_*site*_. The mapping harmonizing these coefficients is then:

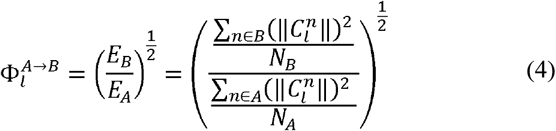

Let *A* be the set of subjects *n* = 1 … *N*_*A*_ scanned with acquisition protocol and scanner defined by site A. We let anything not defined in either expected value to be 1 (resulting in no change when applied). Note that the RISH features come from spherical harmonic coefficients that were normalized to their respective b0 images. As a result, the template has a unitless, intuitive value centered around 1. The template from site B to site A is the inverse of the formula above.

The template for a given order is a single 3-D image volume. To apply 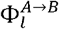, we multiply all spherical harmonic volumes belonging to order *l* by this volume. For example, spherical harmonic volumes 1 through 5 will be multiplied 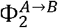 [18].

Conceptually, we are removing the previous target mean rotationally invariant signal and replacing it with that of our reference site.

### B. MidRISH

We find the geometric midpoint in RISH feature between each site. The template then becomes 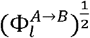. Both sites require template scaling. The inverse, 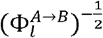, is applied to site B.

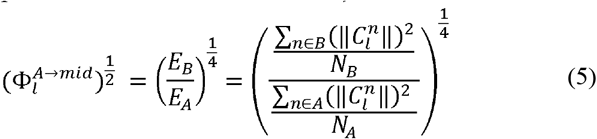

### C. Analysis

We report harmonization efficacy as the difference in site means for 73 regions of interest (cerebral white matter, and 72 white-matter bundles). We compare effect sizes of MD and FA for four harmonization schemes: no harmonization, LinearRISH harmonization with Site B as reference, LinearRISH harmonization with Site A as reference, and MidRISH.

### D. Data

To validate MidRISH at harmonizing data from the same study but different scanners, we analyzed 74 Alzheimer’s Disease Neuroimaging Initiative (ADNI) (adni.loni.usc.edu) patients split between GE 3T and SIEMENS 3T scanners, matched for sex (22 females from each) and age (72.7±6.7 years). Both had b-values of 1000 s/mm^2^ and were sampled at 48 directions and stored with 2 x 2 x 2 mm^3^ resolution. This cohort was launched in 2003 as a public-private partnership, led by Principal Investigator Michael W. Weiner, MD [23].

For the cross-study analysis, we used sex-age-diagnosis matched data from Vanderbilt Memory and Aging Project (VMAP) and Baltimore Longitudinal Study of Aging (BLSA). We considered 117 participants free of cognitive impairment (total of 234) with 65 females and ages 71.9±7.43. Both sites used single shell acquisitions but varied gradient schemes. VMAP acquired 32 directions at a b-value of 1000 s/mm^2^, and BLSA acquired 64 samples at a b-value of 700 s/mm^2^, respectively. BLSA used a Philips 3T scanner at a resolution of 2.2 x 2.2 x 2.2 mm^3^ and resampled to 0.81 x 0.81 x 2.2 mm^3^. VMAP used a Philips 3T scanner at a resolution of 2 x 2 x 2 mm^3^.

### E. DTI and tractometry metrics

From spherical harmonic models, we can resample to reconstruct diffusion signal. We then fit a tensor to this signal and calculate MD and FA. We calculate the mean over all voxels in cerebral white matter defined by the SLANT atlas, a deep whole brain high resolution brain segmentation [24]. To get a better idea of the MidRISH efficacy and LinearRISH bias across white matter, we break this region into 72 bundles defined by the tractseg toolbox (version 2.6)[25]. Tractseg creates a segmentation of bundles based on the DWI, generates tract orientation maps [26] and performs probabilistic tractography with 2000 streamlines in each segment [27]. We extract bundle-wise tractometry (MD and FA) using scilpy (version 1.5.0) [28], [29].

### F. Implementation

#### 1) Code

This code is released on github: https://github.com/nancynewlin-masi/MidRISH.

#### 2) Sensitivity to b-value mapping

While this work performs b-value mapping on BLSA (b=700 s/mm^2^) to project it to VMAP (1000 s/mm^2^). Initial work was limited to sites with similar acquisition parameters (same resolution and b-values) [30], [31]. Later, Karayumak et al proposed b-value mapping for b-values within acceptable range prior to spherical harmonic estimation [32]. It was then implemented for sites with b-values of 900, 1000, and 1300 s/mm^2^ to investigate white matter abnormalities in schizophrenia patients [8]. In this work we provide a comparison between the traditional LinearRISH (Section A) and MidRISH (Section B) using b-value mapping detailed in Section D.3 and without.

#### 3) B-value mapping

LinearRISH and MidRISH methods assume data was acquired at similar gradient strengths. To use these methods when there is a discrepancy in b-value, an additional preprocessing step is needed. Previous work [33] illustrated that diffusion signal has linear decay for values 500 < b-value < 1500 s/mm^2^. The equation for the new diffusion signal 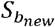 is as follows:

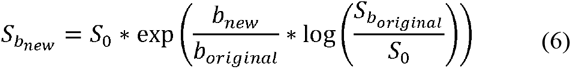

where 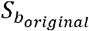 is the original signal collected at 700 s/mm^2^ and *S*_0_ the original baseline. For all analyses we apply this correction before template creation or application when applicable.

#### 4) Diffeomorphic Registration

We create the RISH template with the same protocol for LinearRISH and MidRISH. The resulting template is a voxel-wise scaling factor that is aligned across all subjects in a site and across sites. Thus, we need to register all RISH images to a common space. We used MNI non-linear symmetric template as the registration reference image [34], [35]. Using ANTs, we computed the rigid 3-D registration between each subject’s b0 image and their T1-weighted image, then the non-rigid transformation from T1-weighted image to the reference MNI image [25].

## III. Results

### A. Harmonizing data from different scanners

The difference in MD and FA between the two ADNI sites was not significant and therefore minimal correction was needed. LinearRISH and MidRISH mean MD and FA values are recorded in Table 1. All harmonization schemes produce data closer to the target for MD, however we note that for FA LinearRISH and MidRISH move site means further apart.

**Table 1.**
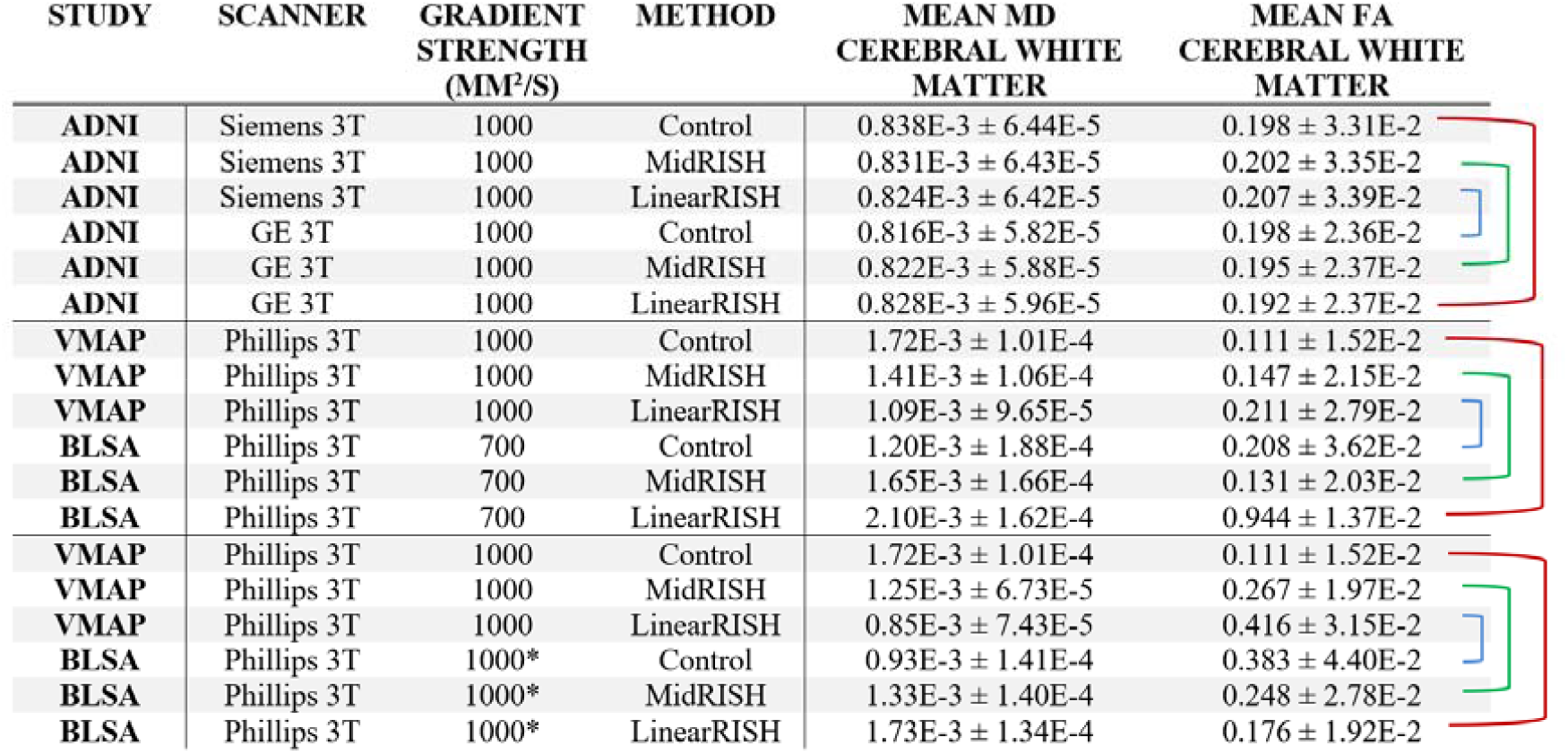
LinearRISH harmonization can be applied to either site DWI and this choice will have downstream effects on DTI derived metrics (MD, FA). Applying LinearRISH to site A shifts mean MD and FA closer to site B’s effect size (blue), and applying LinearRISH to site B shifts mean MD and FA closer to site A’s effect size (red). MidRISH (green) shifts both toward their counterpart site. The middle section reports that harmonizing sites with different b-values causes the projection to overshoot the reference site and incorrectly calibrate the mid-space. This supports the need for b-value correction during preprocessing, indicated by (*) next to the b-values in the third section.

| STUDY | SCANNER | GRADIENT STRENGTH (MM<sup>2</sup>/S) | METHOD | MEAN MD CEREBRAL WHITE MATTER | MEAN FA CEREBRAL WHITE MATTER |
| --- | --- | --- | --- | --- | --- |
| ADNI | Siemens 3T | 1000 | Control | 0.838E-3 ± 6.44E-5 | 0.198 ± 3.31E-2 |
| ADNI | Siemens 3T | 1000 | MidRISH | 0.831E-3 ± 6.43E-5 | 0.202 ± 3.35E-2 |
| ADNI | Siemens 3T | 1000 | LinearRISH | 0.824E-3 ± 6.42E-5 | 0.207 ± 3.39E-2 |
| ADNI | GE 3T | 1000 | Control | 0.816E-3 ± 5.82E-5 | 0.198 ± 2.36E-2 |
| ADNI | GE 3T | 1000 | MidRISH | 0.822E-3 ± 5.88E-5 | 0.195 ± 2.37E-2 |
| ADNI | GE 3T | 1000 | LinearRISH | 0.828E-3 ± 5.96E-5 | 0.192 ± 2.37E-2 |
| VMAP | Phillips 3T | 1000 | Control | 1.72E-3 ± 1.01E-4 | 0.111 ± 1.52E-2 |
| VMAP | Phillips 3T | 1000 | MidRISH | 1.41E-3 ± 1.06E-4 | 0.147 ± 2.15E-2 |
| VMAP | Phillips 3T | 1000 | LinearRISH | 1.09E-3 ± 9.65E-5 | 0.211 ± 2.79E-2 |
| BLSA | Phillips 3T | 700 | Control | 1.20E-3 ± 1.88E-4 | 0.208 ± 3.62E-2 |
| BLSA | Phillips 3T | 700 | MidRISH | 1.65E-3 ± 1.66E-4 | 0.131 ± 2.03E-2 |
| BLSA | Phillips 3T | 700 | LinearRISH | 2.10E-3 ± 1.62E-4 | 0.944 ± 1.37E-2 |
| VMAP | Phillips 3T | 1000 | Control | 1.72E-3 ± 1.01E-4 | 0.111 ± 1.52E-2 |
| VMAP | Phillips 3T | 1000 | MidRISH | 1.25E-3 ± 6.73E-5 | 0.267 ± 1.97E-2 |
| VMAP | Phillips 3T | 1000 | LinearRISH | 0.85E-3 ± 7.43E-5 | 0.416 ± 3.15E-2 |
| BLSA | Phillips 3T | 1000* | Control | 0.93E-3 ± 1.41E-4 | 0.383 ± 4.40E-2 |
| BLSA | Phillips 3T | 1000* | MidRISH | 1.33E-3 ± 1.40E-4 | 0.248 ± 2.78E-2 |
| BLSA | Phillips 3T | 1000* | LinearRISH | 1.73E-3 ± 1.34E-4 | 0.176 ± 1.92E-2 |

### B. Harmonizing data from different studies

MidRISH maintains harmonization efficacy – it has averaged performance between each LinearRISH implementation (Figure 3). Harmonization decreases the difference in mean MD and FA by an order magnitude. This finding is reproduced with mean FA within 72 bundles in Figure 4.

**Fig. 2.**
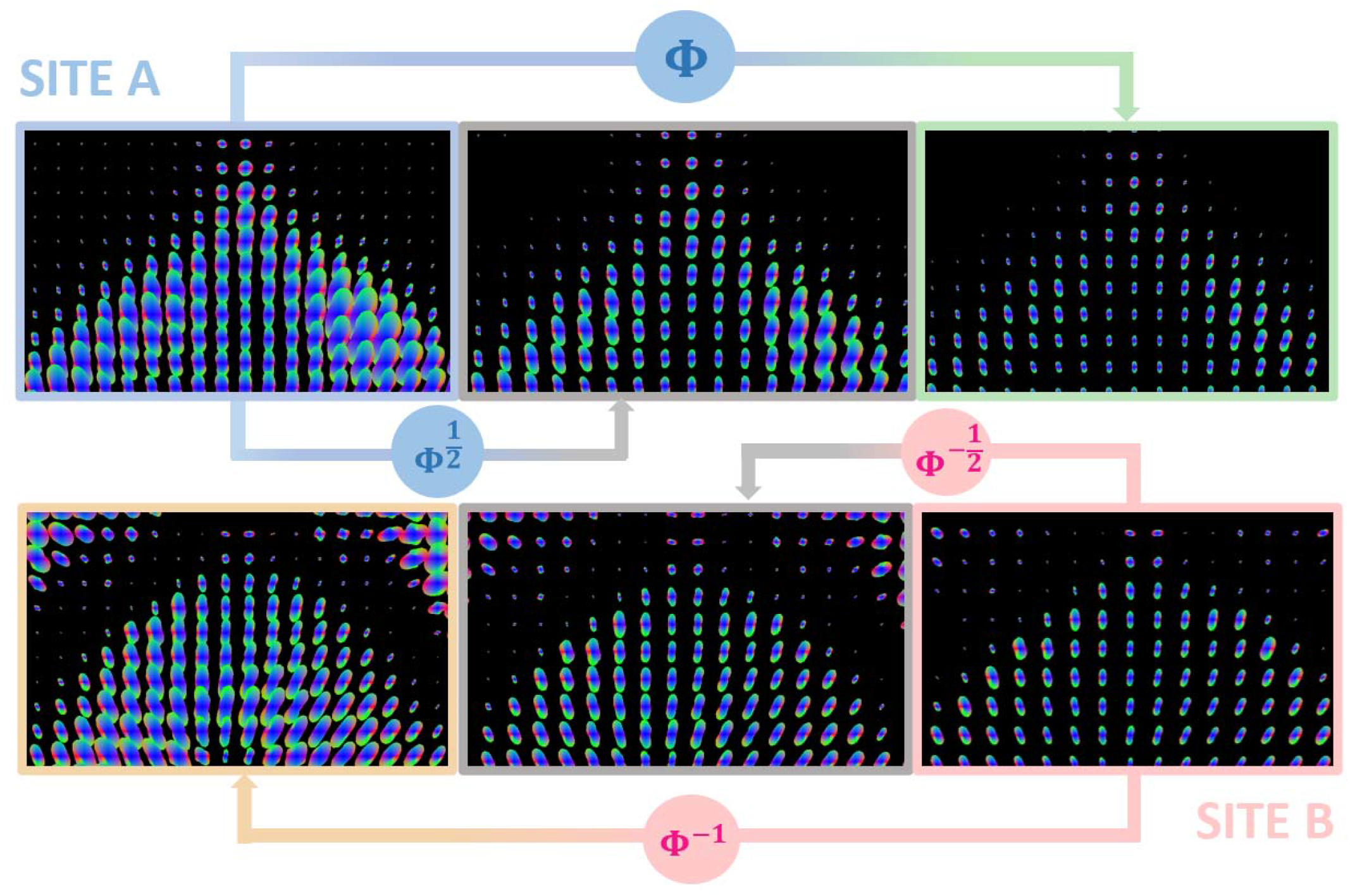
Traditional LinearRISH applies a template Φ or Φ^−1^ to the spherical harmonic representation to harmonize between sites. MidRISH factors in 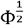 to project each site to a middle space that compromises the bias between sites. Visualizing the 4th order spherical harmonic representation of diffusion signal from the corpus collosum splenium, we see the results of LinearRISH and MidRISH scaling. Note here we are visualizing the signal instead of the orthogonal representation such as DTI.

**Fig. 3.**
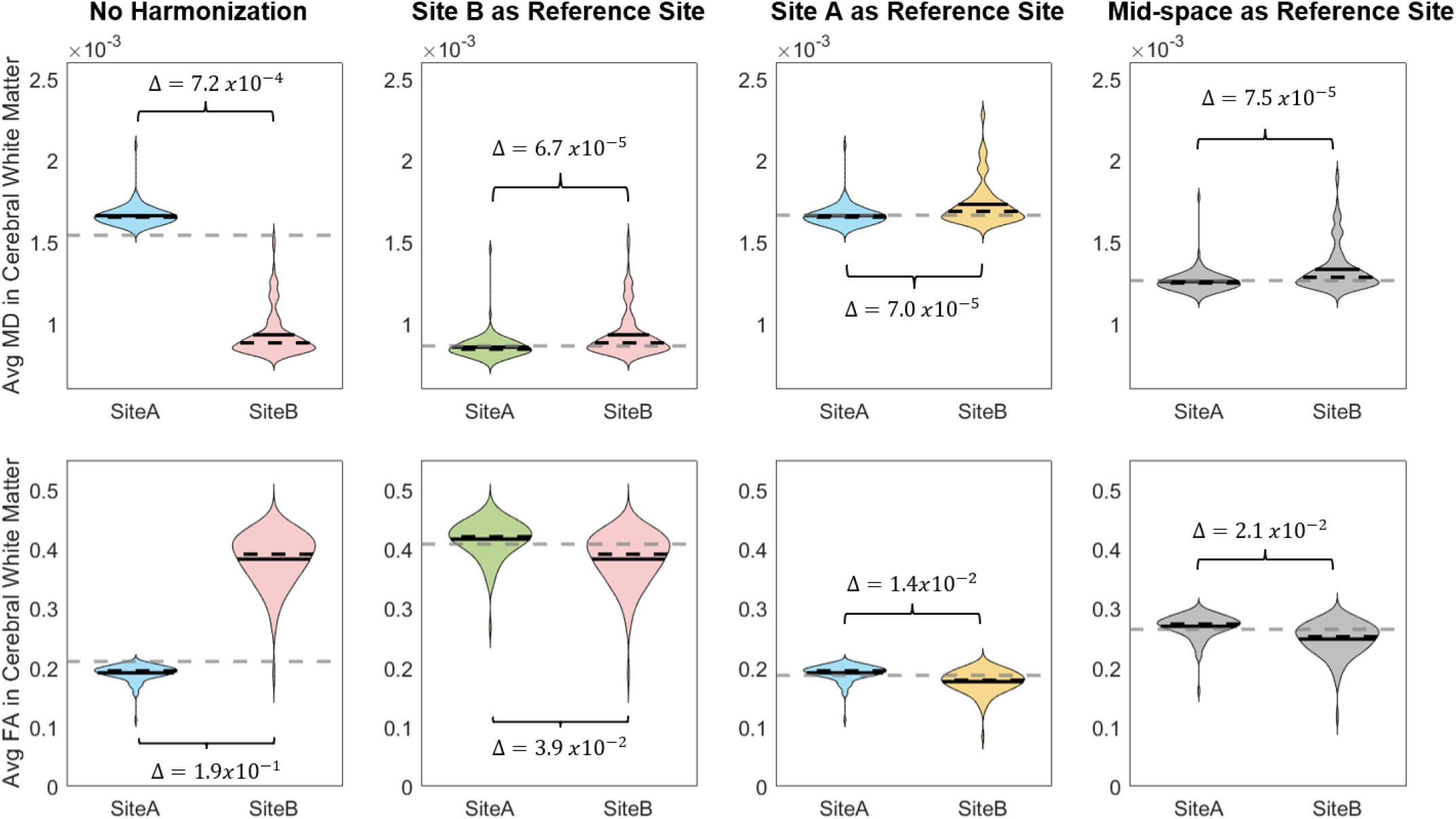
We find significant differences between in MD and FA from Site A, VMAP, and Site B, BLSA (column 1) illustrated with 117 healthy subjects from each site. The difference in site means, Δ, decreases by an order magnitude for both LinearRISH and MidRISH across MD and FA data. However, using BLSA as reference in column 2 causes mean MD of VMAP to decrease and mean FA to increase. Using VMAP as reference in column 3, the mean MD of BLSA increases and mean FA decreases. Clearly, both cannot be quantitatively correct. The proposed approach known as MidRISH provodes a mid-space as reference and projects both sites to their geometric mean. The MidRISH harmonization space results in an average diffusivity and anisotropy across the subject spaces.

**Fig. 4.**
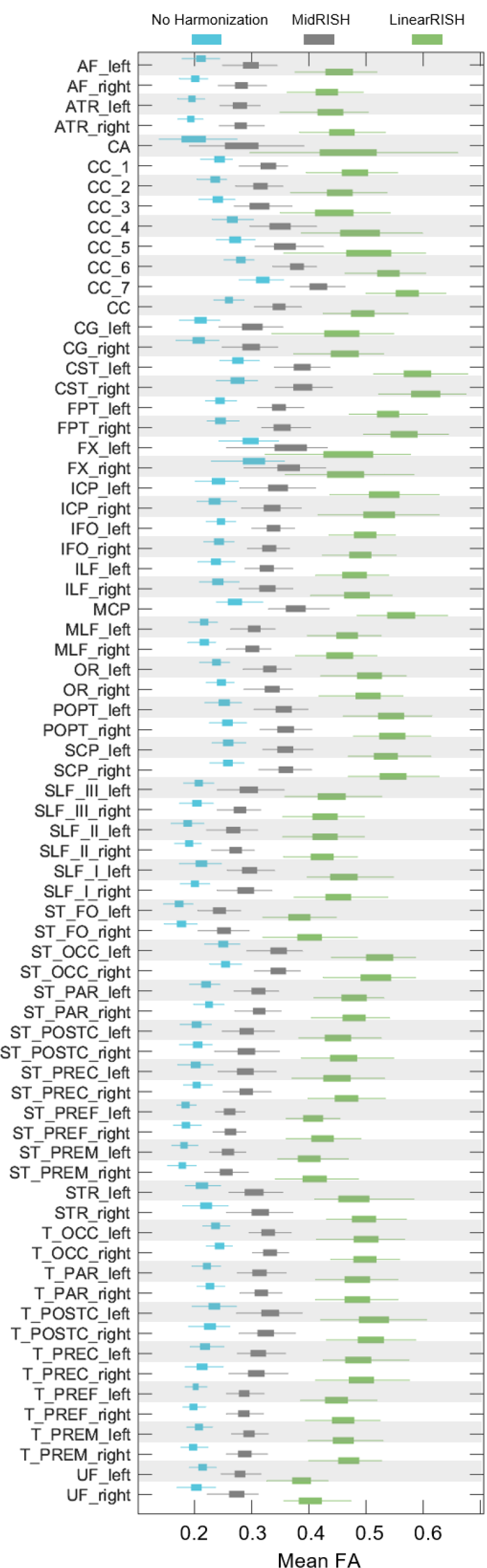
Across 72 anatomically defined bundles, MidRISH (grey) consistently generates tractometry metrics in the middle of the original (blue) and LinearRISH (green) means. For each bundle, we calculated average FA across 117 VMAP scans for each harmonization scheme.

The magnitude of the shift caused by LinearRISH is greater than previously published effect size differences due to multiple sclerosis [9], [36], and Alzheimer’s disease [37]. Inconsistent reference site designation using LinearRISH may result in confounding differences in analysis on pathological DWI studies.

### C. B-value mapping

Results for mean MD and FA with and without b-value mapping are recorded in Table 1. Without prior correction, MidRISH and LinearRISH shift BLSA and VMAP further than the intended MD. We offer this comparison to elucidate that b-value mapping is necessary for effective RISH calculations.

## IV. Discussion and Future Research

With the traditional method, LinearRISH removes site bias in signal by pushing that bias to a user specified parameter, namely the choice of reference site. As we have shown, MidRISH removes this free parameter while maintaining harmonization efficacy. Other limitations of LinearRISH and RISH features in general are still present in MidRISH. For both methods, the key step of template creation is computing the average RISH feature across subjects in each voxel. This calculation necessitates construction voxel-to-voxel registration across subjects to reference space, then across both sites. RISH calculations that follow use this method are sensitive to poor registrations, especially at tissue boundaries.

Extending our choice of midpoint further, we can derive a multivariate midpoint equation for multiple sites. This is

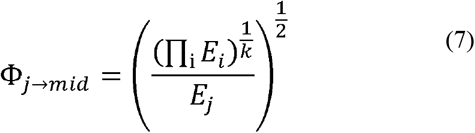

However, many other choices of metric are also reasonable, and we cannot reasonably test or even enumerate all of them. Still, we believe that broadly most choices of midpoint will be better than harmonizing to an arbitrary choice of site.

One limitation of this study is our cohorts were comprised of cognitively unimpaired individuals. As such, we have not accounted for pathologies and their interactions with site variables. While we believe that for most conditions this harmonization will perform adequately, we have only examined normatively healthy control subjects in this paper. Moreover, for extreme anatomy warping conditions (e.g., stroke), harmonization may not be possible at a voxel level.

## V. Conclusion

MidRISH reduces bias of reference selection while preserving harmonization efficacy of traditional LinearRISH. In this paper we illustrate the bias of selection site on effect-size and offer an alternative method that eliminates this step in favor of the geometric mean between sites.

